# Mitochondria drive microglial NRLP3 inflammasome activation via TSPO

**DOI:** 10.1101/2024.03.18.585507

**Authors:** Aarti Singh, Manuel Rigon, Marta Gramaça Caldeira, Danilo Faccenda, Dong Xia, Jordi Lopez-Tremoleda, Zahra Falah Hassan Al-Khateeb, Tong Guo, Rosella Abeti, Paola Giunti, Michelangelo Campanella

## Abstract

Uncontrolled microglial response is core to neuroinflammatory brain diseases. The correlation between the mitochondrial protein TSPO and inflammation has so far failed to explain whether TSPO positively or negatively regulates microglial function. The recent evidence on the species specificity of TSPO in microglia demands a deeper understanding of the protein biology in these brain-resident macrophages. To this end, we have here enrolled a murine model of microglial cells showing that TSPO is required for the priming of mitochondria to inflammation and a conduit for its escalation. Namely, in response to inflammatory cues TSPO is stabilised on the mitochondria where it binds and sequesters NOD-like receptor (NLR) protein (i), represses the PARK2-mediated mitophagy (ii) and engages the retrograde communication with the nucleus via the accumulation of the Nf-kB to promote the expression of pro-inflammatory genes (iii). Notably, the TSPO sustained inflammatory response drives cellular demise and ultimately leads to excitotoxicity (iv).

Our findings advance the current knowledge of TSPO widening the understanding of mitochondria in inflammation and indicating a target for their regulation.

## Introduction

Microglia are the CNS resident immune cells which respond rapidly to external stimuli by switching from a “resting state” to a surveillance mode, to various states of the activation [1]. This evolution is accompanied by a radical cellular morphological change, release of a specific subset of cytokines and subsequent regulation of target proteins, causing either pro-inflammatory (e.g. IL-1β, IL-6, IL-12, TNFα, CD86 and iNOS) [2, 3] or anti-inflammatory (e.g. IL-4, IL-10, IL-13, TGF-β and CD206) downstream effects [4].

A prerequisite for microglial activation is the intracellular accumulation and assembly of the nucleotide oligomerization domain (NOD)-like receptor family pyrin domain-containing 3 (NLRP3) inflammasome. The inflammasome is a complex composed of the NLRP3 protein (i), the adaptor molecule apoptosis-associated speck-like protein containing a caspase recruitment domain (ASC) (ii) and caspase 1 (CASP-1) (iii). The abnormally increased formation of this complex is implicated in the pathogenesis of several chronic pathologies (e.g. Alzheimer’s disease [AD], Parkinson’s disease [PD], multiple sclerosis [MS] and amyotrophic lateral sclerosis [ALS]).

Upon its assembly, the inflammasome triggers the autocleavage of the Caspase-1 protein, activating it. Once cleaved and active, CASP-1 can then process the proinflammatory cytokines pro-IL-1β and pro-IL-18 into their biologically active mature forms [5], enhancing this way inflammation and leading to an inflammatory pyroptotic cell death.

Studies have demonstrated that mitochondria are crucial for NLRP3 inflammasome activation [5, 6]. Mitochondrial damage induced by NLRP3 activators mediates the externalization of mitochondria-derived molecules, including cardiolipin and mt-DNA. This event allows the mitochondrial recruitment and assembly of NLRP3, leading to an inflammasome activation [7, 8]. Consequently, the autophagic removal of damaged mitochondria reduces the NLRP3-mediated inflammatory response [9].

The translocator protein (TSPO) is an evolutionarily conserved five-transmembrane domain protein that localizes to the outer mitochondrial membrane (OMM) [10]. While TSPO is ubiquitously expressed in mammalian systems, the protein is found at high levels in tissues with active lipid metabolism [11]. TSPO is primarily involved in regulating mitochondrial cholesterol influx and steroidogenesis [12]. More recently, biochemical, and pharmacological studies have unveiled that TSPO takes part in a plethora of cell pathways encompassing mitophagy (i), Ca^2+^ signalling (ii) and reactive oxygen species (ROS) generation (iii).

The protein level of TSPO is upregulated in several disease-associated conditions including cancer and central nervous system (CNS) inflammation [13, 14]. TSPO-binding drugs are enrolled on a vast spectrum of therapeutic interventions ranging from anti-neuroinflammatory treatments [15, 16] to chemotherapy [17–19] and anxiolysis [3]. TSPO-ligands, such as the isoquinoline PK11195 have been applied for the diagnostic imaging of brain injury, during which increased TSPO expression is associated with neuroinflammatory damage. Targeting of TSPO is therefore used to measure brain inflammation both in experimental models and human patients [20, 21].

Recent findings suggest that reduced autophagic competence promotes pro-inflammatory microglial polarisation [22]. Taken together, the above evidence raised the speculation that TSPO is involved in inflammasome signalling pathways as TSPO is indicated to counteract mitochondrial autophagy. However, the molecular details, especially the interplay between TSPO, mitochondria and NLRP3 inflammasome activation, are still elusive. Therefore, it remains difficult to elaborate on the role of TSPO during microglial activation.

In the current study, we demonstrate that TSPO mediates inflammasome signalling in microglia cells. Along with the repression of mitophagy, TSPO sustains the scaling of the inflammatory response in microglial cells via the stabilization of the NLRP3 on the mitochondria and exploitation of the retrograde signalling. A complex scaling of inflammation which irreversibly drives microglial cells to death.

## Material and Methods

### Cell Culture and Transfection

BV2 microglial cells (ATCC) were maintained in a temperature-controlled, humidified incubator at 37 °C with 5 % CO_2_ (Hera Cell 240, Thermo Scientific, Essex, UK). TSPO KO microglia were generated using GeneArt™ CRISPR Nuclease Kit (Invitrogen, A21174) and maintained like the Wild Type (WT) BV2. The cells were cultured in Dulbecco’s Modified Eagle Medium (DMEM) containing High Glucose (25 mM), L-Glutamine (4 mM) and Sodium Pyruvate (1 mM) (Thermo Fisher, 11995065), supplemented with 10% Fetal Bovine Serum (FBS) (Thermo Fisher, 10082147) and 1% of 100 U/mL Penicillin and 100 mg/mL streptomycin (Thermo Fisher, 15140122). GE-180 was a gist by General Electric Healthcare. Transfection was performed using Lipofectamine 3000 transfection reagent (Thermo Fisher, L3000015) according to the manufacturer’s instructions and following optimization.

### Immunocytochemistry (ICC)

Cells were seeded onto Nunc™ Lab-Tek™ II Chamber Slide™ System (Thermo Fisher, 154534PK) treated and/or transfected as described. After treatment, cells were washed twice using 0.01 M Phosphate Buffered Saline (PBS 1X) and fixed using 4% paraformaldehyde (PFA) solution for 15min at RT. After 3X washes with PBS permeabilization was made with 0.3% Tritonx-100 for 15min. The cells were subsequently washed with PBS (3X) for 10-minute incubation periods. The slides were blocked for 1h in 5% FBS (blocking buffer) at RT and incubated overnight at 4^D^C in a humidified chamber after the addition of primary antibody diluted in blocking buffer. The slides were then washed 3x15mins using PBS-T (PBS with 0.2% Tween) and then incubated with secondary antibody in PBS-T for 1-2h at RT. The secondary antibodies were conjugated to Alexa Fluor dyes (Life Technologies, UK). Following washing (3 x 15 minutes using PBS-T) in the dark slides were mounted using 4’, 6-diamidino-2-phenylindole (DAPI) mounting solution (Abcam, ab104139).

### Immunohistochemistry (IHC)

For the IHC a total of 8 adult, 10-week-old male mice, weighing 30-34 g (Charles River Laboratories, Margate, UK), were used. Mice were housed in an Individually Ventilated Cage (IVC; Allentown Europe, UK), in a 12 h light-dark cycle, with controlled room temperature (21±1°C) and relative humidity (40-60%) and with diet and water ad libitum. All animal procedures were carried out under a Project Licence approved by the Animal Welfare and Ethical Review Body, at Queen Mary University of London and the UK Home Office, by the EU Directive 2010/63/EU. Animals were subjected to controlled cortical impact (CCI) injury. Briefly, animals were acclimatised for 1- week after arrival to the animal unit. Mice were anaesthetized (ketamine 50 mg/kg and medetomidine 0.5 mg/kg; IP) and placed in a stereotaxic frame. A right lateral craniotomy was carried out, 2.0 mm behind bregma and 2.5 mm lateral to the midline and a CCI injury was induced using a 3 mm impactor tip with a speed of 3 m/s, a depth of 2.2 mm and a dwell time of 100 ms, applied with the PCI3000 Precision Cortical Impactor™ (Hatteras Instruments, Inc., US). After the injury, the skull flap was placed back unfixed to allow for expansion, and the skin was sutured. Buprenorphine (0.05 mg/kg, s.c.) was used in all animals pre-operatively for pre-emptive analgesia and post-operatively every 12 h for 3 days post-surgery. Naïve animals were used as controls. Slides were scanned at x40 using the In Cell Analyser 2200 (INCA2200) System (Cytiva, Marlborough, United States) and the In Cell Developer Toolbox v1.9.2 (Cytiva, Marlborough, United States) was used to quantify the signal. Image processing was done using ImageJ software (National Institutes of Health, Bethesda, USA).

### RT-qPCR

RNeasy Plus mini kit (Qiagen, 74134) was employed to extract the RNA according to the manufacturer’s instructions and then quantified using a NanoDrop spectrophotometer (ThermoFisher Scientific). Quantinova Reverse Transcription Kit (Qiagen, 205413) was used according to the manufacturer’s instructions to transcribe the RNA to cDNA. The 2X Quantinova SYBR green master mix (Qiagen, 204141). The qPCR reaction was performed in 384 well plates in triplicate reactions. The mean Ct was normalized to the Ct for a housekeeping gene glyceraldehyde-3-phosphate dehydrogenase (GAPDH). The delta Ct (ΔCt) was calculated: gene of interest Ct – housekeeping gene Ct. From this, the relative mRNA content is calculated using the following formulae: 2^^-ΔΔ^Ct. The mean 2^^-ΔΔ^Ct was calculated from the 4-5 independent experiments. Sequences of Primers are reported in **Table 1**.

### Immunoblotting

Protein concentrations were quantified using the detergent-compatible (DC) protein assay (Bio-Rad, 5000112) according to manufacturer instructions. The absorbance was subsequently read using a plate reader (Tecan Infinite M200 Pro, UK) at 695-750 nm with a BSA standard curve used to determine the protein concentration (μg/μL) of the samples. The volume of protein required to achieve 20 μg of sample was loaded onto 12% polyacrylamide gels. Bio-Rad Mini-PROTEAN tetra system electrophoresis unit was used. After transfer the blots were blocked in a 5 %(w/v) solution of milk powder in TBS-T (50 mM Tris, 150 mM NaCl, 0.05% Tween 20 (Sigma, P9416)) for 1h at RT. The membrane was incubated with antibodies diluted in milk at 4°C overnight. Following primary antibody incubation, the membranes were washed 3x5min with a 5 % (w/v) solution of milk powder in TBS-T at RT. The membrane was incubated for 2 h with the corresponding peroxidase-conjugated secondary antibody diluted in milk. Membranes were washed (3x5min) in TBS-T. To visualize the blot the ECL western blotting detection kit (Amersham, RPN2133) was utilized. Imaging and visualization were performed using a ChemiDoc™ MP System (Bio-Rad, 1708280).

### Co-Immunoprecipitation

Microglia BV2 WT cells and BV2 TSPO KO cells were seeded in 6-well plates and treated with LPS 100 ng/ml for 24 hours or LPS 100 ng/ml for 24 hours + 2.5 mM ATP for 30 min. After treatment, the cells were washed twice in PBS and subsequentially lysed with standard Lysis buffer (150 mM NaCl, 1% Triton in 50 mM Tris-HCL pH 8) and protein concentrations were determined via DC protein assay as described above and 60 μg of total protein from this sample have been loaded in the input (INP) control lanes for the Western Blot. For the immunoprecipitation assay, 1 μg of TSPO antibody (Abcam, ab108489) has been added to 50 μl of magnetic beads linked to Protein A (ThermoFisher Scientific, 10001D) for 15 minutes at RT to induce the binding. After the incubation, the beads were washed twice with PBS and incubated overnight with 100 ug of total protein from the samples obtained above. The next day, the Flow Through (FT) fraction was collected and the beads were washed twice with PBS. Finally, the proteins attached to the beads and the TSPO antibody have been eluted by adding 60 μl of Loading Buffer (375 mM Trsi-HCL pH 6.8, 6% SDS, 4.8% Glycerol, 9% β-mercaptoethanol, 0.03% bromophenol blue) and boiling the samples for 10 minutes, obtaining this way the IP fraction for the Western Blot analysis.

### Cholesterol assay

The Amplex Red Cholesterol Assay Kit (Thermo Fisher, A12216 was used to quantify cholesterol levels. Cells were resuspended in the 1X reaction buffer (0.1 M potassium phosphate, pH 7.4, 0.05 M NaCl, 5 mM cholic acid, 0.1% Triton X-100) that was supplied with the kit. The suspension was aliquoted in triplicate in 96-well plates. Amplex Red reagent containing 2 U/mL HRP, 2 U/mL cholesterol oxidase, and 0.2 U/mL cholesterol esterase was prepared according to the kit instructions. The reaction was incubated at 37 °C for 30 minutes (in dark). The subsequent change in fluorescence was measured in a Tecan M200 Pro Plate Fluorescence microplate reader (Tecan, UK) using excitation at 544 nm and emission at 590 nm.

### Reactive Oxygen Species (ROS) analysis

Dihydroethidium (DHE) is sensitive to O^2-^ and oxidises to 2-hydroxyethidium and the rate at which the nuclear signal rises in intensity correlates with cytosolic levels of superoxide. Cells were seeded on 22 mm glass coverslips and treated for 24h either with Vehicle or LPS. 24h post-treatment cells were washed using room temperature PBS 1X and transferred to an Attofluor Cell Chamber (Sigma Aldrich, A7816), 5 μM of DHE was added in HBBS medium (Hanks’ Balanced Salt Solution, Thermo fisher, 24020117) and the increase in intensity measured continuously over time using a UV Nikon microscope.

### Nitric oxide (NO2) levels analysis

Griess reaction allows for the spectrophotometric detection of nitrite formed by the oxidation of NO quantified using a microplate reader at an absorbance of 548 nm. Cells were seeded and treated in triplicate on 96 well plates 48 hours before the experiment. Sample media was used alongside photometric references made using the Griess reagent diluted in deionized water and sodium nitrite standards (1 – 100 μM) and quantified according to the manufacturer’s instructions.

### Cytotoxicity assessment

LDH (lactate dehydrogenase) activity assay (Merck) is a colourimetric assay used to measure the concentration of LDH in the cell culture medium. Microglial BV2 cells were seeded in 25 cm^2^ flasks and either cultured for 48h or cultured for 24h and then treated using LPS for another 24h. The microglial-conditioned medium was collected, filtered, and used to treat neuroblastoma N2a cells for 24h. N2a cells were also treated with complete medium and complete medium supplemented with 5µM glutamate. N2a cells seeded in triplicate in the same plate and treated with complete medium only were used as a negative control, whereas N2a cells seeded in triplicate in the same plate and treated with glutamate were used as a positive cytotoxicity control. The average absorbance values of the triplicate samples were calculated. Cytotoxicity % for each of the conditions was calculated using the formula recommended by the kit company: (experimental value -negative control)/(positive control - negative control)*100. Cytotoxicity % was further normalised to be expressed as a percentage of the WT + Vehicle condition. The experiment was repeated 6 times, with BV2 and N2a cells being seeded on 6 different days. LDH activity was assessed following the manufacturer’s instructions. Sample absorbance was read at 409/492nm with a Tecan Sunrise plate reader.

### Transcriptome data analysis

Total RNA samples were extracted using Quick-RNA Miniprep (Zymo Research, USA) according to the manufacturer’s instructions, and then quantified with an Agilent 2100 bioanalyzer. The quantified RNA samples were used for NEBNext® rRNA-depleted (Human/Mouse/Rat) stranded library preparation. Library preparation and RNA sequencing were conducted at the UCL Genomics. Libraries were prepared using non-strand-specific Illumina TruSeq Sample Preparation Kits followed by Illumina sequencing. FASTQ files were aligned using TopHat and Cufflinks. Normalization and differential analyses were carried out using R software Bioconductor[61] package DESeq2[62], and gene set enrichment analysis (GSEA) was carried out using EGSEA[63]. For gene expression profiling, the lists of genes belonging to NLRP3 inflammasome complex and autophagosome were retrieved with Gene Ontology accession GO:0072559 and GO:0005776 respectively. Genes involved in mitophagy were retrieved from GO:0000423 and Reactome accession R-HSA-5205647.

**Methods for the assessment of Ca^2+^ signalling, Oxygen Consumption Rate (OCR), mitochondrial membrane potential (ΔΨ_m_), cellular viability and GE-180 binding are reported in Supplementary Material**

### Statistical analysis

Three biological and technical replicates have been attained for each of the experiments.

All statistical analyses were performed in GraphPad Prism 8.0 (GraphPad Software, USA). Data are presented as mean ± standard error of the mean (SEM). Variations between three or more independent groups were determined using a one-way analysis of variance (ANOVA). Two-way ANOVA with randomized block was used to evaluate differences between groups while accounting for the day block effect. Tukey’s post hoc test was used to reveal all possible pairs of means within the data sets. P values of less than 0.05 were considered significant (P < 0.05 *; P < 0.01 **; P < 0.001 ***; P < 0.0001 ****).

## Results

### TSPO knockout enhanced the respiration and proliferation of BV2 cells

Our previous data showed that treatment of BV2 cells with lipopolysaccharides (LPS) triggered a significant increase in the expression levels of TSPO protein, indicating a correlation between TSPO and *in vitro* pro-inflammatory activation (M1 polarization) of microglial cells [23]. In the current study, we discovered that the TSPO accumulate in Iba1-enriched microglial cells. following traumatic brain injury in mice. (**Figure 1a, b**). Meanwhile, BV2 cells treated with the anti-inflammatory cytokine interleukin (IL-4) [24], displayed a decreased expression level of TSPO compared to the control (**Figure 1c, d**).

**Figure 1.**
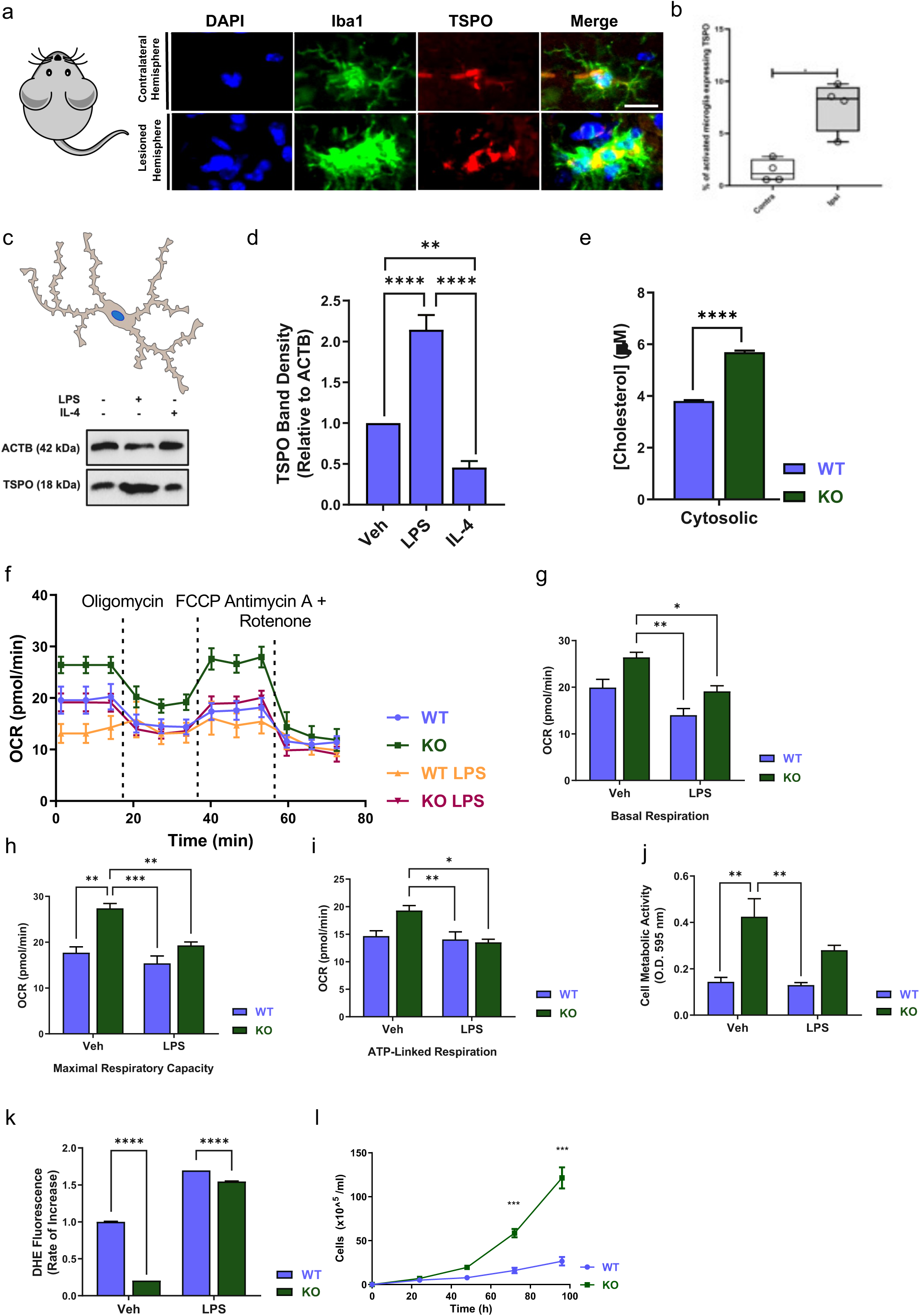
Generation and analysis of TSPO knocked out microglial cells. **a, b**) Immunofluorescent analysis of brain slices from traumatic brain injured mice staining DAPI. Images showing Iba 1 (as a marker of microglial activation) and TSPO labelling (**a**) with relative quantification (**b**) **c**, **d**) The anti-inflammatory cytokine IL-4 does not promote TSPO expression. Western blotting analysis of TSPO protein levels in BV2 cells treated with LPS (100 ng/ml) or IL-4 (20 ng/ml) (**c**) and relative quantification (**d**). **e**) Analysis of intracellular cholesterol levels in WT and TSPO KO BV2 cells. Cells were fractionated and total cholesterol content was quantified in cytosolic fractions. TSPO KO cells have higher levels of cholesterol in the cytosol **f, g, h, i, j**) Bioenergetic profiles of WT and TSPO KO BV2 cells at rest and after 24 h LPS treatment (100 ng/mL). Extracellular flux analysis was carried out using a standard Mito Stress Test kit with 1.5 µM oligomycin, 1 µM FCCP, and 0.5 µM rotenone/antimycin A (**f**). OCR values, expressed as pmol O_2_/min, were used to calculate basal respiration (difference between OCR at rest and after rotenone/antimycin A administration) (**g**), ATP-linked respiration (difference between OCR at rest and upon oligomycin treatment) (**h**) and maximal respiratory capacity (difference between OCR under FCCP treatment and following the addition of rotenone/antimycin A) (**i**). The analysis indicates that TSPO KO cells have a higher metabolic capacity than WT. **j)** MTT viability assay showing increased mitochondrial activity in TSPO KO cells (OD three times higher than WT), which is retained even after treatment with LPS (100 ng/ml, 24 h). **k)** Analysis of intracellular ROS levels in WT and TSPO KO cells using the red fluorescent superoxide indicator DHE. Cells were incubated with either vehicle (Veh) or LPS (100 ng/ml, 24h). TSPO KO BV2 cells have significantly lower levels of ROS when compared to WT cells, either treated or untreated. **l)** Cell proliferation rate comparison between BV2 WT and TSPO KO BV2 cells over 100 hours.

To further elaborate the role of TSPO during M1 polarization, we generated a TSPO- knockout microglial cell model by using the CRISPR/Cas9 technology in BV2 cells (hereafter referred to as TSPO KO) (**Supp. Figure 1**). To characterize the phenotypic changes induced by TSPO knockout, we assessed the intracellular distribution of cholesterol, as TSPO is a well-established intracellular transporter [25]. Our results showed that the cytosolic cholesterol level is significantly higher in the TSPO KO cell line confirming the loss of function caused by TSPO depletion (**Figure 1e**). We next compared the basal mitochondrial respiratory capacity between wild-type and TSPO KO cells, as well as in the presence of LPS. Interestingly, TSPO deletion promotes mitochondrial oxidative phosphorylation, exemplified by increased mitochondrial activity (**Figure 1f-i**), increased metabolic activity (**Figure 1j**), decreased cytosolic superoxide (**Figure 1k)** and a greater cell proliferation rate (**Figure 1i**).

### TSPO ablation blunts pro-inflammatory responses and modulates Ca^2+^ signalling in BV2 cells

We next investigated the effect of TSPO knockout on the pro-inflammatory response capacity of BV2 cells, by looking into the activation of NLR family pyrin domain containing 3 (NLRP3) inflammasome. NLRP3 inflammasome activation requires a two-step mechanism. The first step, namely the priming step, involves activation of the NF-κB pathway, leading to upregulation of pro-IL-1β and NLRP3 protein levels, and the second step is featured by the assembly of NLRP3, ASC and pro-caspase-1 into the NLRP3 inflammasome complex [26].

Immunoblot results showed that ablation of TSPO prevents the increase in NLRP3 protein expression in response to LPS or LPS combined with ATP (**Figure 2a,b**). Immunocytochemistry experiments also showed that NLRP3 accumulates to a lesser extent in the mitochondria in the absence of TSPO (**Figure 2c, d**). qPCR analysis showed that TSPO knockout decreased the transcription of inducible nitric oxide synthase (iNOS), NLRP3 and the pro-inflammatory cytokine IL-1β, (**Supp. Figure 2).** Wild-type cells also displayed a greater mitochondrial network compared to TSPO KO cells (**Figure 2e).** Co-immunoprecipitation assay showed that LPS treatment induced the interaction between NRLP3 and TSPO and such interaction was diminished to the basal level when wild-type cells were treated with LPS in combination with ATP (**Figure 2f**).

**Figure 2.**
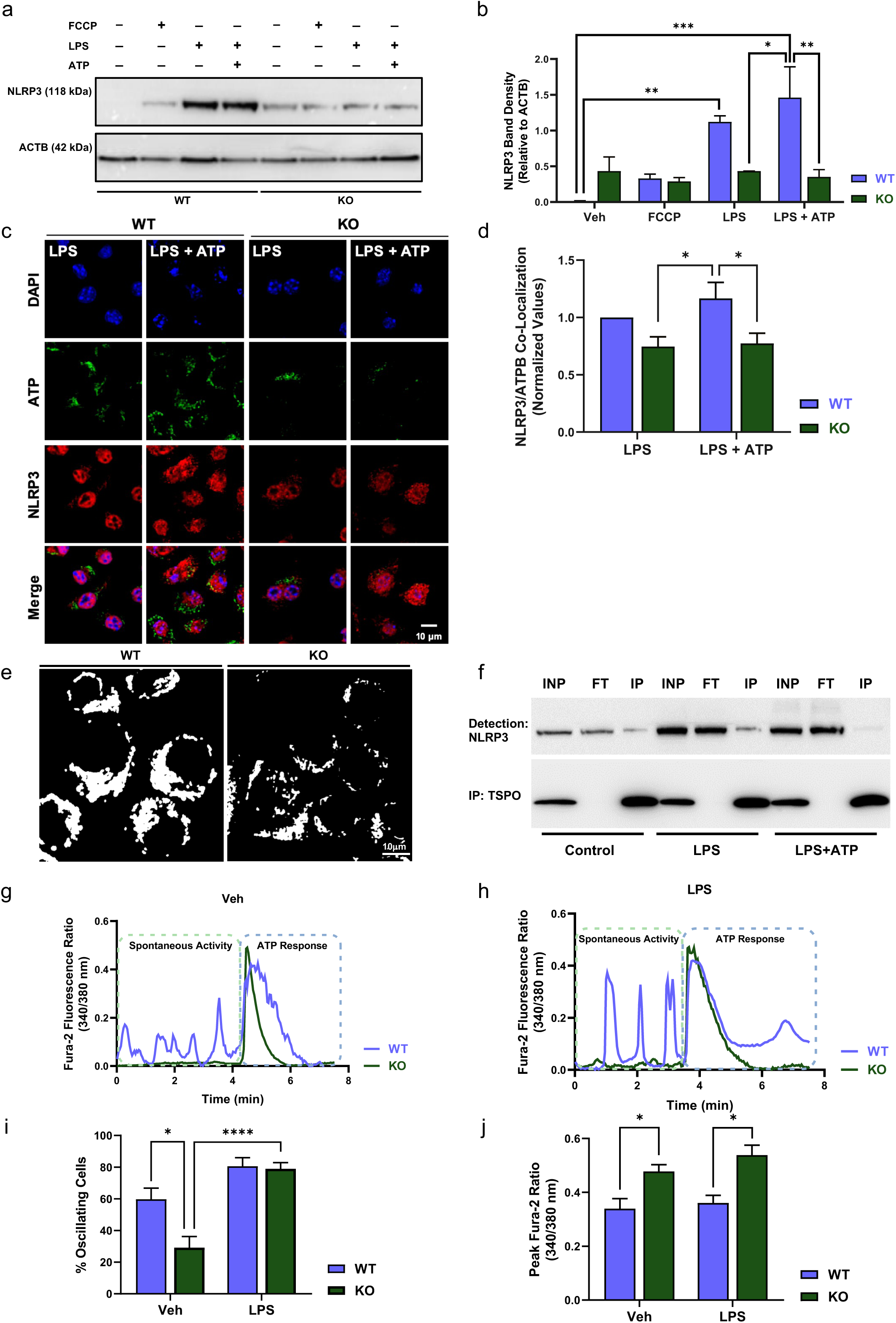
TSPO ablation prevents NRLP3 stabilization on mitochondria and represses Ca^2+^ accumulation in the cytosol. **a**, **b**) Western blotting analysis of NLRP3 protein levels in WT and TSPO KO BV2 cells at rest and after FCCP, LPS or LPS/ATP treatments (as described above). The representative blot (**a**) indicates that both LPS and LPS/ATP-induced activation result in a significant upregulation in NLRP3 protein expression in WT cells only, while NLRP3 levels don’t change when TSPO is ablated, as quantified (**b**). **c, d**) Immunocytochemistry analysis of occupancy of mitochondria by NLRP3 in LPS- and LPS/ATP-activated WT and TSPO KO BV2 cells (as previously described). The representative images (**c**) show BV2 cells stained with anti-NLRP3 (red) and anti-ATP synthase subunit beta (ATPB) (green) antibodies. The quantification of the co-localization between the two fluorescent signals is reported in panel **d**. Knocking out TSPO inhibits pro-inflammatory mitochondrial recruitment of NLRP3. **e)** Mito-tracker staining of mitochondrial network WT and TSPO KO BV2 cells (n=3). **f)** Co-Immunoprecipitation between TSPO and NRLP3 (n=4). **g, h**) Live-cell imaging analysis of intracellular Ca^2+^ dynamics in resting and activated (100 ng/ml LPS, 24 h) BV2 cell lines. Intracellular Ca^2+^ levels were monitored by using the ratiometric, fluorescent Ca^2+^ indicator Fura-2 AM, before and after the addition of ATP (100 µM). Representative traces of the Fura-2 ratio at rest (spontaneous activity) and after ATP administration are specifically reported in untreated (**g**) and LPS-treated cells (**h**). **i)** Quantification of the percentage of oscillating cells in BV2 WT and TSPO KO cell lines in resting conditions or activated by LPS after the addition of ATP. **l**) Quantification of the intracellular Ca^2+^ peak response in the conditions described above.

Activation of the NLRP3 inflammasome complex and the activation of microglial cells is accompanied by intracellular Ca^2+^ signalling [27–29]. Whether TSPO knockout modifies the intracellular Ca^2+^ signalling was investigated. Intracellular Ca^2+^ levels were measured by the ratiometric, fluorescent Ca^2+^ indicator Fura-2 AM, and Ca^2+^ mobilization induced by the addition of ATP. As shown in **Figure 2g**, before stimulation with ATP, WT BV2 cells presented spontaneous Ca^2+^ oscillatory activity that was greatly reduced in TSPO KO cells (**Figure 2i**). Treatment with LPS intensified these oscillations which nonetheless remained lower in TSPO KO cells (**Figure 2h**). Administration of ATP elicited a rapid release of Ca^2+^ in both untreated and LPS-treated cells, but the TSPO KO BV2 cells showed a slightly higher amplitude in cytosolic Ca^2+^ transient and faster clearance compared to their WT counterpart (**Figure 2j**).

### TSPO knockout induced upregulation of mitophagy and autophagy in BV2 cells

RNA-Seq analysis unveiled a global picture of the changes in the transcriptome of BV2 cells caused by TSPO knockout. Out of the genes which displayed responsiveness to TSPO knockout and LPS treatment, we focused on the executors of the NRLP3 inflammasome in the pro-inflammatory pathways and the autophagy/mitophagy regulators (**Figure 3a, Supp. Figure 3a**). In general, TSPO knockout induced the downregulation of members of the NRLP3 inflammasome and upregulated the genes related to autophagy and mitophagy (**Figure 3a)**. In line with the RNA-Seq result, immunoblot data demonstrated that key makers of autophagy and mitophagy are elevated in TSPO KO cells at both the resting condition and upon LPS treatment, including i) LC3 lipidation (**Figure 3b, c**), ii) mitochondrial translocation of the p62 (SQSTM1) (**Figure 3d, e**), iii) PARK2 accumulation (**Figure 3f, g**), and, iv) extensive mitochondrial ubiquitination with MTCO1 degradation (**Figure 3h-j**).

**Figure 3.**
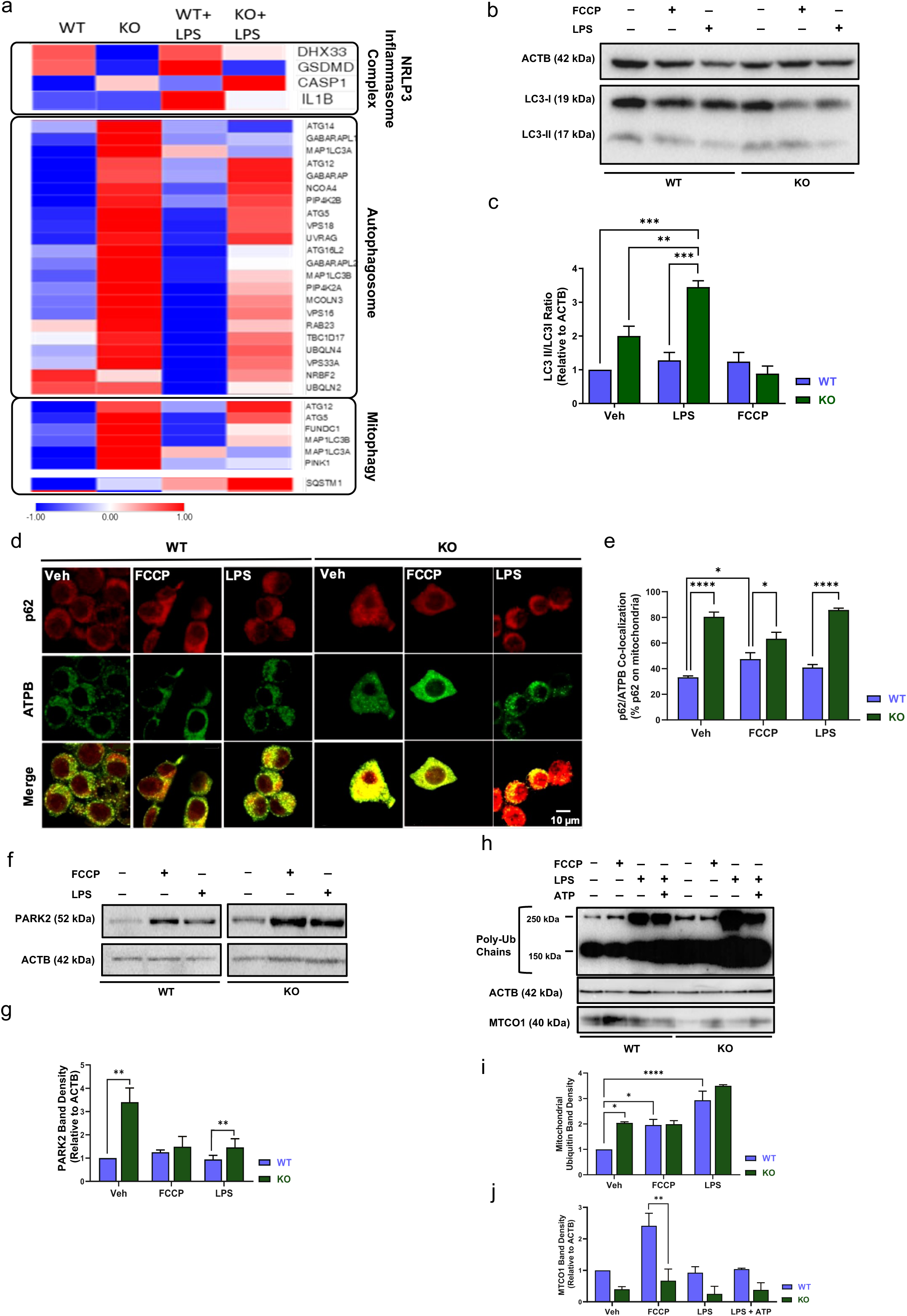
TSPO represses macro and targeted autophagy during microglial cell activation. **a)** Heat maps of the RNA-seq profiles in BV2 WT and BV2 TSPO KO cells in resting condition and after LPS treatment (100 ng/mL, 24 h). **b**, **c**) Measurement of LC3 lipidation (LC3-II formation) in WT and TSPO KO BV2 cells following 24 h treatment with LPS (100 ng/mL, 24 h) or FCCP (10 uM, 24 h). Both LC3 bands (**b**) were quantified and the LC3-II: LC3-I ratio of expression was calculated (**c**) to compare LC3 lipidation (LC3-II formation) between samples before and after treatment. TSPO KO cells have a higher LC3-II: LC3-I ratio both at rest and following LPS-induced activation, suggesting that the autophagic flux increases in the absence of TSPO. **d, e**) Representative co-immunocytochemical analyses of SQSTM1/P62 and ATPB in BV2 WT and BV2 TSPO KO microglial cells untreated, exposed to LPS or LPS+ATP (**d**). **e**) Histogram reporting the relative quantification. **f**, **g**) Representative immunoblot (**f**) and densitometry analysis (**g**) of mitochondrial Parkin levels in WT and TSPO KO BV2 cells at rest and following treatment with either FCCP (10 µM, 24 h), LPS (100 ng/mL, 24 h) or LPS (100 ng/mL, 24 h)/ATP (2.5 mM, 30 min). TSPO KO cells are characterized by increased basal expression of Parkin. **h**, **i, j**) Western blotting analysis of MTCO1 protein levels and mitochondrial ubiquitination levels in WT and TSPO KO BV2 cells at rest or after treatment with either FCCP, LPS or LPS/ATP (same as above). As presented in panel **h** and quantified in the panel **I**, TSPO KO cells have an overall decreased expression of MTCO1, the level of which does not change following treatments. MTCO1 expression is instead greatly induced by FCCP in WT cells. **j**) Quantification of the mitochondrial ubiquitination levels

### TSPO is required for the LPS-induced nuclear retro-translocation of NF-kB

Results described above suggest absence of TSPO downregulates the NLRP3 inflammasome activation. Among all the events during the activation cascades, ASC is affected to a lesser extent by TSPO knockout, compared to NLRP3 and IL-1β **(Figure 2 and Supp. Figure 2).** We herein reason that the priming step of NLRP3 inflammasome activation is more closely related to TSPO. In addition, recently, it was revealed that mitophagy activation is linked to NF-kB activation but is independent of the caspase-1 activation [22]. Since our results described above indicate TSPO ablation attenuates inflammasome activation whereas induced mitophagy in BV2 cells, we, therefore, speculate that such attenuation is mediated by NF-κB. Both immunocytochemistry and immunoblot confirmed that in TSPO KO cells, there is a significantly decreased level of mitochondrial and cytosolic NF-κB (**Figure 4a-d**), whereas an elevated level of nuclear NF-κB. Correspondingly, TSPO ablation changed the secretome content of BV2 cells upon LPS treatment. Wild-type and TSPO KO BV2 cells were treated with either vehicle control or LPS for 24 h. The microglial conditioned media (Mic-CM) medium of each condition is collected and transferred to neuroblastoma N2a cells with for 24h. Following incubation, neuronal cytotoxicity is assessed by the lactate dehydrogenase (LDH) assay (**Figure 4g, h**). Mic-CM from LPS-treated wild-type BV2 cells induced more cell death of N2a cells compared to that from LPS-treated TSPO KO cells, indicating that loss of TSPO reduced microglia-induced neurotoxicity, potentially through suppressing the release of pro-inflammatory cytokines [22]. Additionally, TSPO KO cells showed less sensitivity to LPS-induced cell death (**Figure 4i**) suggesting that TSPO ablation reduces pro-inflammatory type of cell death (i.e. pyroptosis). Ligands of the protein were then tested as markers and modulators of inflammation.

**Figure 4.**
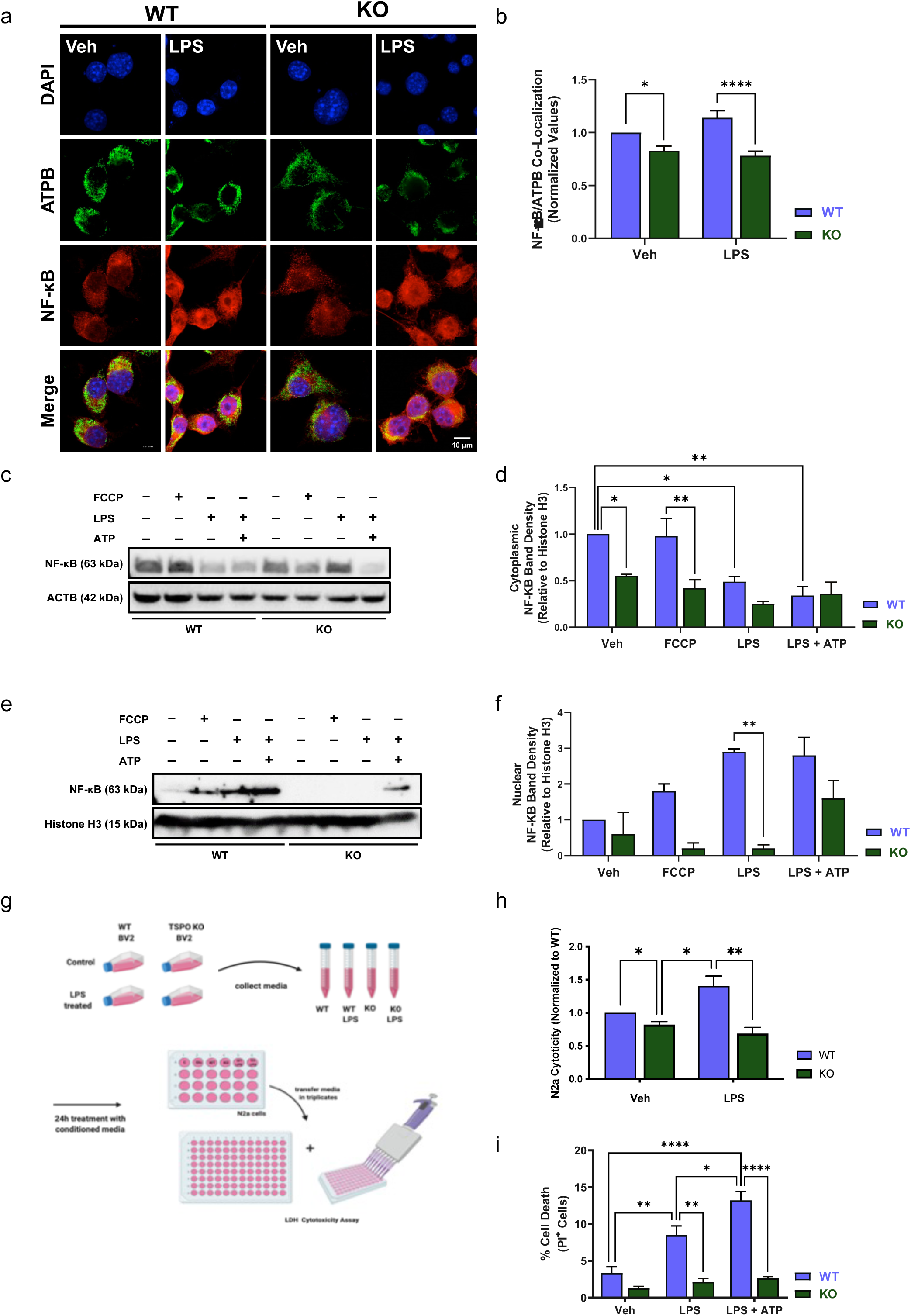
TSPO engages NF-kB in microglial pathogenic signalling. **a**, **b**) Fluorescence ICC analysis of LPS-induced NF-κB nuclear translocation in WT and TSPO KO BV2. Cells were treated with 100 ng/mL LPS for 24 h. Representative images (**a**) showing NF-κB immunostaining (red) in combination with DAPI nuclear counterstain (blue). The levels of NF-κB nuclear fluorescence, reported in panel **b**, indicate that TSPO ablation inhibits LPS-mediated nuclear translocation of NF-κB. **c**-**f**) Analysis of NF-κB cytosolic and nuclear translocation in WT and TSPO KO BV2 cells following pro-inflammatory stimuli. Cells were treated with either FCCP (10 µM, 4 h), LPS (100 ng/mL, 24 h) or LPS/ATP (100 ng/mL LPS for 24 h followed by 2.5 mM ATP for 30 min). Cytosolic and nuclear fractions were then used to monitor the relocation of NF-κB between the two compartments via Western blotting. The representative blots (**c**, **e**) and respective densitometry analyses (**d**, **f**) show that treatment with LPS or LPS/ATP causes a significant translocation of NF-κB into the nucleus in WT cells, which is hampered by the lack of TSPO. **g**, **h**) Analysis of microglia-induced neurotoxicity in mouse N2a cells exposed for 24 h to Mic-CM media from either untreated or LPS-treated (100 ng/ml, 24 h) WT and TSPO KO BV2 cells. WT BV2 cell Mic-CM media induced neuronal cytotoxicity, which was measured through quantification (**h**) of LDH released in the extracellular environment. N2a cells treated with TSPO KO cell Mic-CM did not show signs of cytotoxicity. **i)** Quantification of changes in cell death as measured by PI inclusion between WT and KO cells treated with LPS (24 hours using 100 ng/mL) or LPS ATP (24 hours using 100 ng/mL followed by 30 min using 2.5 mM ATP) to induce pyroptosis. TSPO KO protects from LPS-induced pyroptosis.

### TSPO is a target for the pharmacological control of inflammatory response

Previous results have endorsed the potential of TSPO as an *in vivo* biomarker of mitochondrial pathology for PET imaging [30]. To further validate TSPO as a viable pharmacological target to reduce pathological neuroinflammatory conditions, we tested the effect of a recently developed, highly specific TSPO ligand, namely ^18^F- GE-180 (S-N, N-diethyl-9-[2-18F-fluoroethyl]-5-methoxy 2,3,4,9-tetrahydro-1H- carbazole-4-carboxamide). ^18^F-GE-180 (hereafter referred to as GE-180) is the lead compound from a new series of tricyclic indoles, which have been shown to have a high affinity for TSPO. GE-180 has been developed as a PET tracer for *in vivo* imaging of TSPO (**Figure 5a-c**). ^18^F-GE-180 repressed the production of CD68 protein as well as nitrite ion, which are two indicators of active microglia (**Figure 5d- f**). In the absence of LPS, GE-180 didn’t alter the basal level of CD68 and nitrite ions. In contrast, wild-type cells when treated with GE-180 displayed a decreased responsiveness level to LPS, whereas no significant difference was detected in TSPO KO cells, indicating GE-180 attenuated TSPO-mediated microglial activation. In support of this finding, GE-180 reduced the NF-kB retro-translocation caused by LPS treatment in wild-type cells (**Figure 5g, h**). Taken together, our data demonstrated that TSPO is a key mediator of microglia pro-inflammatory activation, whose function can be specifically counteracted by GE-180.

**Figure 5.**
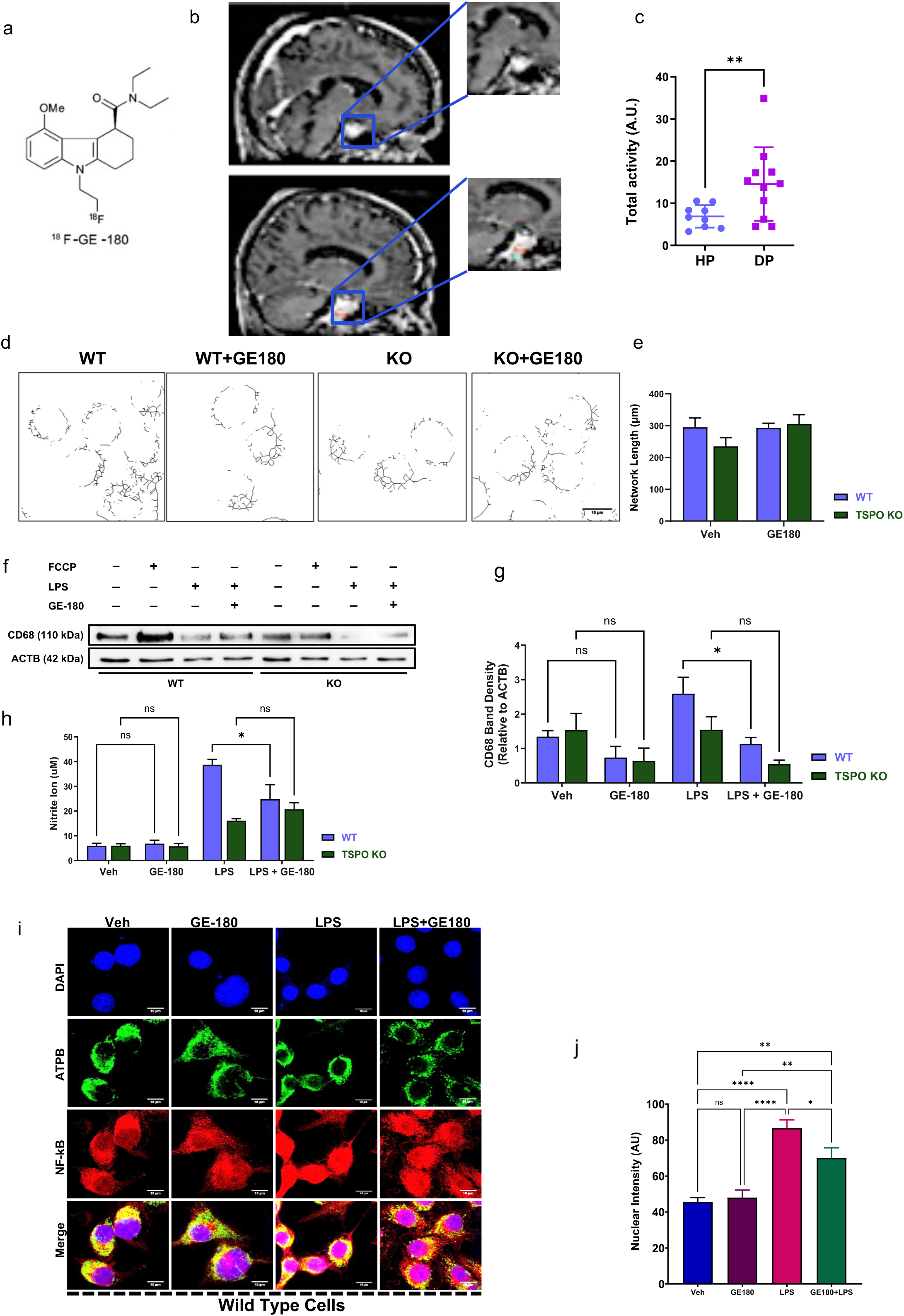
TSPO is a pharmacological target to prevent pro-inflammatory microglial cell activation and associated demise. **a-c**) Molecular structure and improved standard uptake values (SUV) of the binding of novel TSPO ligand, GE-180 with the chemical structure shown in (**a**) and representative PET-magnetic resonance imaging (MRI) scans with and without PMOD-generated “masks” on the carotid (**b**) with the quantification demonstrating significant upregulation in binding of ligand in the diseased patient (DP) over healthy patient (HP) (**c**). [18F]GE-180 activity was evaluated around maximal TSPO increase. Total disease activity measured for both groups of patients (HP: healthy patients; DP: diseased patients) normalized to standard uptake value (SUV) measured in carotid arteries (highlighted in multicolour rectangles at the bottom). **d)** Representative skeletonized images of mitochondrial morphology read via green mito-tracker in BV2 cells treated with GE-180 and relative analysis reported in **e**. **f, g**) Immunoblotting analysis of CD68, Actin B and TSPO levels expression in WT and TSPO KO BV2 cells treated with LPS (100 ng/mL, 24h), or GE-180 (100nM, 24h) with quantification in (**g**) that demonstrates a significant downregulation in CD68 accumulation in KO cells after LPS treatment. **h)** Griess reagent assay displays the changes in nitrite levels between WT and TSPO KO cells treated with LPS (100 ng/mL, 24h), or GE-180 (100nM, 24h). The panel reiterates the reduction in the profile of neuroinflammatory markers in KO cells even after LPS administration. **i)** Immunocytochemical data on NF-kB retro-translocation on the nucleus in response to LPS (100 ng/ml, 24 h) alone or combined with GE-180 (100nM, 24h) in WT and TSPO KO cells. Nuclear intensity measured via co-localization is reported in **j**.

**Figure 6.**
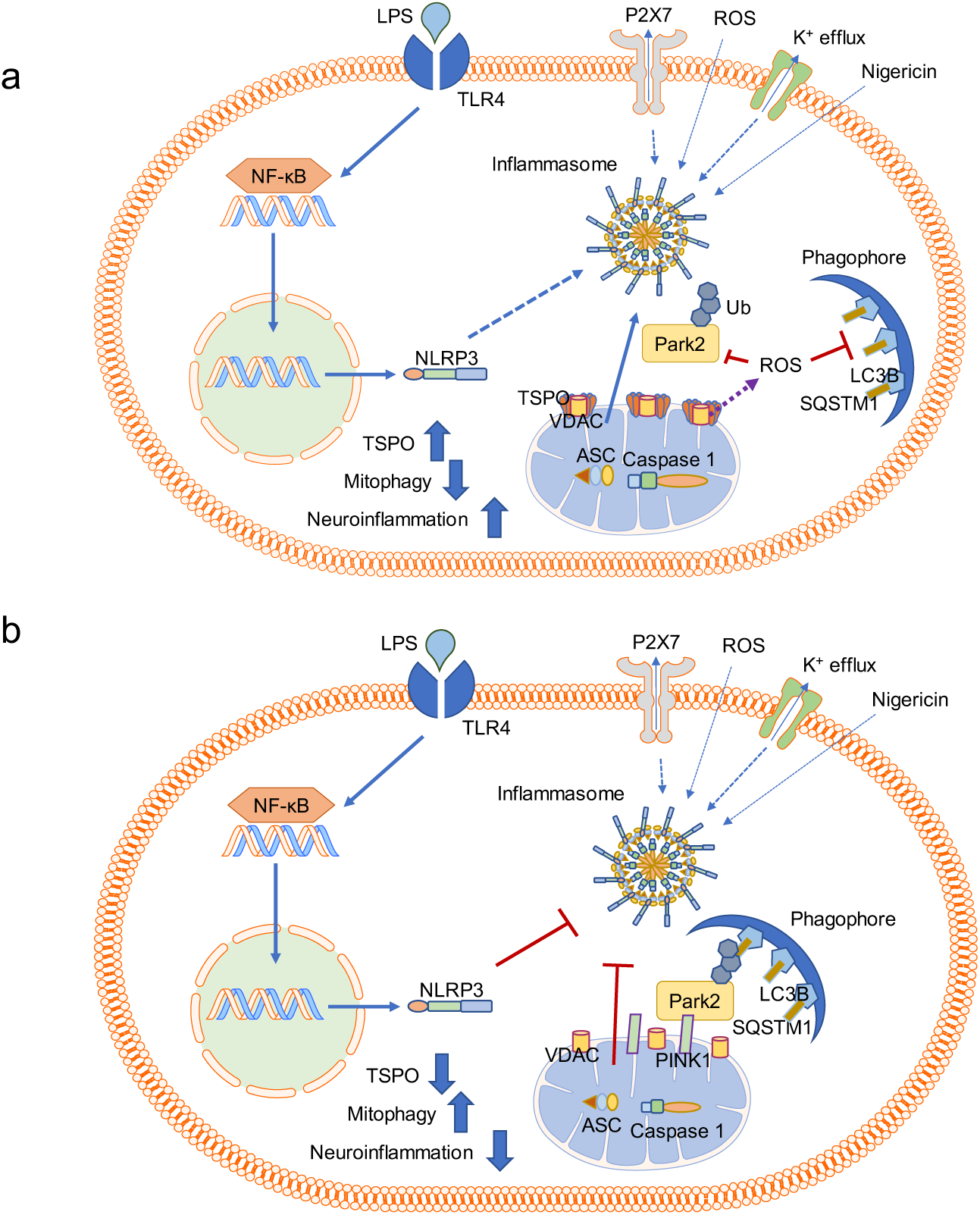
Working Model. **a, b)** Models of NLRP3 inflammasome assembly and activation according to TSPO expression. Following a priming step consisting of inflammatory TLR engagement, NF-κB gets activated and induces NLRP3 and IL-1β expression; inflammasome assembly is driven by a second activating stimulus, commonly a danger signal increasing intracellular ROS, K^+^ efflux or extracellular ATP, which leads to caspase- 1-mediated IL-1β and IL-18 secretion and then pyroptosis. Lack of TSPO promotes autophagic clearance of mitochondria, thereby impeding mitochondrial recruitment of NLRP3 and inflammasome assembly. Hence, TSPO KO microglial cells have reduced expression of pro-inflammatory IL-1β and iNOS after priming by the Toll-like receptor (TLR) agonist LPS.

We propose a new model of TSPO regulation of neuroinflammation based on our data. In this model, we propose that the NLRP3 inflammasome is activated in a TMPO-dependent manner. Removal of TSPO resulted in attenuation in nuclear recruitment of NF-κB, leading to reduced NLRP3 inflammasome activation, and reduced production of pro-inflammatory cytokines. On the other hand, since NF-kB can function as an inhibitor of mitophagy and autophagy, the lack of active nuclear NF-kB due to TSPO depletion gave rise to autophagy and mitophagy. Our data highlighted an essential role for TSPO as a regulator of the inflammatory response of microglia cells by fine-tuning the mitophagy pathway and the assembly of the NLRP3 inflammasome.

## Discussion

Cumulatively, these findings reveal significant phenotypic changes in microglial cell lines according to TSPO on which translocation and activation of the NRLP3 inflammasome is dependent.

Previously we learnt that TSPO drives cellular survival to stress by facilitating the mitochondrial retro-communication with the nucleus via the formation of contact between the two organelles [31]. Here, we show that TSPO primes microglial cells to their inflammatory response ensuing the production of pro-inflammatory mediators which are instead lost in TSPO KO cells (**Figure 2 a-d; Supp. Figure 1 c-e**).

Reduction in mitochondrial respiratory capacity and oxidative metabolism are prodromic to this (**Figure 1h-k**) acknowledging a loss in homeostasis which is mirrored by the higher rate of redox stress (i) (**Figure 1m**) and the impaired Ca^2+^ homeostasis (ii) (**Figure 2i, j**; **Supp. Figure 2c-e**). TSPO reduces the intracellular storage capacity for Ca^2+^ leading to fluctuations of the cation in the cytosol also at resting conditions. Such a modification in the core cellular signalling seems therefore to place the TSPO “competent” microglia into a pro-active state which is highly reactive to LPS which not only promotes the redistribution of NLRP3 on the mitochondria (**Figure 2 c, d)** but also the binding with TSPO (**Figure 2 h**).

Such an upscaling of the inflammatory response is easily traceable in the transcriptome data which highlights the prominent inflammatory phenotype in microglial cells expressing TSPO (**Figure 3a, b**).

Though the mitochondrial translocation of NLRP3 was previously demonstrated [35, 36], the mechanic of such redistribution is here for the first time linked to a stress-induced protein. TSPO has been for years acknowledged as a pristine target in brain inflammation [32, 33] for which there are ligands capable of anti-inflammatory function [16, 34, 35]. Strictly linked to this the TSPO ligand Ro5-4864 was found to attenuate NLRP3 activation [36] by preventing mitochondrial damage and reducing superoxide accumulation.

The reduction of TSPO expression is beneficial in various inflammatory and neurodegenerative conditions [37–41] as well as the removal of mitochondria via autophagy (mitophagy) a suitable mechanism to counteract inflammation [42].

It is therefore not surprising that TSPO negatively regulates mitophagy in microglia (**Figure 3 d-i**) like it does in other models of analysis [43–47] causing accumulation of ROS [44, 48]. Reducing the expression of TSPO is therefore capable of increasing mitophagy and reducing ROS (**Figures 1** and **3**). Hence, we speculate that TSPO-mediated counteraction of mitophagy, by making the mitochondrial network bigger, offers a platform for the rapid execution of the inflammatory signalling via the stabilization of the NRLP3 inflammasome complex.

The positive correlation between NLRP3 and TSPO has been previously reported in patients with bipolar disorders [46] whose mononuclear cells present an increased production of TSPO (i), paralleled by an increase in the NLRP3 inflammasome complex-forming proteins and a reduction in those associated with mitophagy (iii) [46].

Previously it was also postulated that mitophagy coincides with a decrease in several inflammatory molecules, meaning that mitophagy regulation dictates the degree of inflammation and vice versa [22, 49–51]. Equally, it was shown how NF-kB transcriptionally controls p62/SQSTM1 to prevent overexploitation of inflammation [22]. We have therefore enquired about such a signalling pattern in our model of study discovering that NF-kB accumulation in the nucleus is positively regulated by TSPO (**Figure 4 a-c**).

Furthermore, TSPO is required for the execution of the NLRP3-Inflammasome-Caspase-1 pathway triggered by LPS thus representing a limiting step for the inflammatory type of cell death known as pyroptosis [52]. Knocking out of TSPO is therefore protective from cellular demise and an equal effect is attained with the ligand of the protein GE-180 (**Figure 5 a-c**) which, capable of restraining the nuclear accumulation of Nf-kB, prevents the activation of the inflammatory proteins (**Figure 5 f-j**) without modifying the mitochondrial network (**Figure 5 d,e**).

This suggests an effect beyond mitophagy and thus TSPO overexpression *per se* does not completely prevent the recycling of mitochondria, suggesting that a threshold level might exist for which cells adapt to the inflammatory environment compromising between accumulation and clearance of damaged mitochondria.

It is potentially within this threshold that it might stand the beneficial physiological effect of TSPO in engaging the NLRP3 inflammasome assembly correctly and instead the pathological exploitation of NRLP3 amplification hence the consequent demise due to caspase-1 engagement.

Notably, recent findings suggest activation of TSPO expression is accompanied by microglial activation in mice and rats, but such a correlation is currently absent in non-human primate disease models or human brains with common neurodegenerative and neuroinflammatory diseases [53]. Given that BV2 cells are microglial cells derived from C57/BL6 murine, this manuscript mechanistically advances our understanding of TSPO’s pro-inflammatory role in this species whilst offering means to further elucidate the differences reported in the human counterpart.

## Acknowledgements

The research activities led by M.C. are supported by the following funders, who are gratefully acknowledged: Biotechnology and Biological Sciences Research Council [grant numbers BB/M010384/1 and BB/N007042/1; The European Research Council COG 2018 - 819600_FIRM; Fondation ARC pour la Recherche sur le Cancer ARCLEADER2022020004901; AIRC-MFAG 21903, Rotary Foundation, GiME Charity and MIUR (PRIN-2022 Prot. 202252ZLSX).

An iCase Studentship awarded by the BBSRC and co-funded by General Electric Healthcare has supported A.S. in her endeavours.

We would like to thank Dr Andrew Hibbert for the help with the imaging data, and Dr Ruby Chan for the assistance in the statistical analysis.

## Authors Contribution

M.C. conceived, designed, and coordinated the project together with A.S. who has performed almost the entirety of the in-cell experiments, their analyses and curation including the data points attained with GE-180. J. L.T. and Z. F. H. A. have run the work on animal tissues. M.R. conducted the biochemical assessment of mitochondrial ubiquitination, assisted A.S. in the autophagy/mitophagy experiments and conducted the co-immunoprecipitation analysis, its analysis and presentation. The data on intracellular Ca^2+^signalling which were conceived by M.C., D.F, and A.S. were conducted by D.F. and M.G.C. who assisted with their design too. M.G.C. finalised the assay on N2a cytotoxicity designed together with M.C. R.A., P.G. and D.X. performed the RNA-seq and their analysis. A.S. wrote the manuscript which was edited by M.C. and finalised in the current format with the key assistance of T.G. and M.R. All authors have critically reviewed and approved the manuscript.

## References

1. Ousman, S.S. and P. Kubes, Immune surveillance in the central nervous system. Nature Neuroscience, 2012. 15(8): p. 1096–1101.

2. Smith, J.A., et al., Role of pro-inflammatory cytokines released from microglia in neurodegenerative diseases. Brain Res Bull, 2012. 87(1): p. 10–20.

3. Wang, D.-s., et al., Anxiolytic-like effects of translocator protein (TSPO) ligand ZBD-2 in an animal model of chronic pain. Molecular Pain, 2015. 11(1): p. 16.

4. Cherry, J.D., J.A. Olschowka, and M.K. O’Banion, Neuroinflammation and M2 microglia: the good, the bad, and the inflamed. J Neuroinflammation, 2014. 11: p. 98.

5. Gurung, P., J.R. Lukens, and T.D. Kanneganti, Mitochondria: diversity in the regulation of the NLRP3 inflammasome. Trends Mol Med, 2015. 21(3): p. 193–201.

6. Zhou, R., et al., A role for mitochondria in NLRP3 inflammasome activation. Nature, 2011. 469(7329): p. 221–5.

7. Booshehri, L.M. and H.M. Hoffman, CAPS and NLRP3. J Clin Immunol, 2019. 39(3): p. 277–286.

8. Zhong, Z., et al., New mitochondrial DNA synthesis enables NLRP3 inflammasome activation. Nature, 2018. 560(7717): p. 198–203.

9. Kim, M.J., et al., SESN2/sestrin2 suppresses sepsis by inducing mitophagy and inhibiting NLRP3 activation in macrophages. Autophagy, 2016. 12(8): p. 1272–91.

10. Bonsack, F. and S. Sukumari-Ramesh, TSPO: An Evolutionarily Conserved Protein with Elusive Functions. Int J Mol Sci, 2018. 19(6).

11. De Souza, E.B., et al., Peripheral-type benzodiazepine receptors in endocrine organs: autoradiographic localization in rat pituitary, adrenal, and testis. Endocrinology, 1985. 116(2): p. 567–73.

12. Rone, M.B., J. Fan, and V. Papadopoulos, Cholesterol transport in steroid biosynthesis: role of protein-protein interactions and implications in disease states. Biochim Biophys Acta, 2009. 1791(7): p. 646–58.

13. Ory, D., et al., PET radioligands for in vivo visualization of neuroinflammation. Curr Pharm Des, 2014. 20(37): p. 5897–913.

14. Rupprecht, R., et al., Translocator protein (18 kDa) (TSPO) as a therapeutic target for neurological and psychiatric disorders. Nat Rev Drug Discov, 2010. 9(12): p. 971–88.

15. Mages, K., et al., The agonistic TSPO ligand XBD173 attenuates the glial response thereby protecting inner retinal neurons in a murine model of retinal ischemia. Journal of Neuroinflammation, 2019. 16(1): p. 43.

16. Scholz, R., et al., Targeting translocator protein (18 kDa) (TSPO) dampens pro-inflammatory microglia reactivity in the retina and protects from degeneration. Journal of Neuroinflammation, 2015. 12(1): p. 201.

17. Jia, J.B., et al., Novel TSPO-targeted Doxorubicin Prodrug for Colorectal Carcinoma Cells. Anticancer Res, 2020. 40(10): p. 5371–5378.

18. Mendonça-Torres, M.C. and S.S. Roberts, The translocator protein (TSPO) ligand PK11195 induces apoptosis and cell cycle arrest and sensitizes to chemotherapy treatment in pre-and post-relapse neuroblastoma cell lines. Cancer Biol Ther, 2013. 14(4): p. 319–26.

19. Takai, N., et al., PK11195 enhances chemosensitivity to cisplatin and paclitaxel in human endometrial and ovarian cancer cells. Int J Mol Med, 2010. 25(1): p. 97–103.

20. Crawshaw, A.A. and N.P. Robertson, The role of TSPO PET in assessing neuroinflammation. J Neurol, 2017. 264(8): p. 1825–1827.

21. Zhang, L., et al., Recent developments on PET radiotracers for TSPO and their applications in neuroimaging. Acta Pharmaceutica Sinica B, 2021. 11(2): p. 373–393.

22. Zhong, Z., et al., *NF-*κ*B Restricts Inflammasome Activation via Elimination of Damaged Mitochondria*. Cell, 2016. 164(5): p. 896–910.

23. Wright, S.D., et al., CD14, a Receptor for Complexes of Lipopolysaccharide (LPS) and LPS Binding Protein. Science, 1990. 249(4975): p. 1431–1433.

24. Murray, P.J., Macrophage Polarization. Annual Review of Physiology, 2017. 79(1): p. 541–566.

25. Owen, D.R., et al., TSPO mutations in rats and a human polymorphism impair the rate of steroid synthesis. Biochemical Journal, 2017. 474(23): p. 3985–3999.

26. Bauernfeind, F.G., et al., *Cutting Edge: NF-*κ*B Activating Pattern Recognition and Cytokine Receptors License NLRP3 Inflammasome Activation by Regulating NLRP3 Expression*. The Journal of Immunology, 2009. 183(2): p. 787–791.

27. Eichhoff, G., B. Brawek, and O. Garaschuk, Microglial calcium signal acts as a rapid sensor of single neuron damage in vivo. Biochimica et Biophysica Acta (BBA) - Molecular Cell Research, 2011. 1813(5): p. 1014–1024.

28. Koizumi, S., et al., UDP acting at P2Y6 receptors is a mediator of microglial phagocytosis. Nature, 2007. 446(7139): p. 1091–1095.

29. Wang, D.-s., et al., Anxiolytic-Like Effects of Translocator Protein (TSPO) Ligand ZBD-2 in an Animal Model of Chronic Pain. Molecular Pain, 2015. 11: p. s12990–0013.

30. van den Ameele, J., et al., [(11)C]PK11195-PET Brain Imaging of the Mitochondrial Translocator Protein in Mitochondrial Disease. Neurology, 2021. 96(22): p. e2761–e2773.

31. Desai, R., et al., Mitochondria form contact sites with the nucleus to couple prosurvival retrograde response. Sci Adv, 2020. 6(51).

32. Bonsack, F., C.H. Alleyne, Jr., and S. Sukumari-Ramesh, Resveratrol Attenuates Neurodegeneration and Improves Neurological Outcomes after Intracerebral Hemorrhage in Mice. Front Cell Neurosci, 2017. 11: p. 228.

33. Dou, X., et al., *[Down-regulation of TSPO expression doesn’t affect the productions of TNF-*α, *IL-1*β *and IL-6 in LPS-stimulated BV-2 microglia].* Xi Bao Yu Fen Zi Mian Yi Xue Za Zhi, 2014. 30(9): p. 897–900.

34. Azrad, M., et al., The TSPO Ligands 2-Cl-MGV-1, MGV-1, and PK11195 Differentially Suppress the Inflammatory Response of BV-2 Microglial Cell to LPS. Int J Mol Sci, 2019. 20(3).

35. Bae, K.R., et al., Translocator protein 18 kDa negatively regulates inflammation in microglia. J Neuroimmune Pharmacol, 2014. 9(3): p. 424–37.

36. Lee, J.W., et al., A translocator protein 18 kDa ligand, Ro5-4864, inhibits ATP-induced NLRP3 inflammasome activation. Biochem Biophys Res Commun, 2016. 474(3): p. 587–593.

37. Campbell, G. and D.J. Mahad, Mitochondrial dysfunction and axon degeneration in progressive multiple sclerosis. FEBS Lett, 2018. 592(7): p. 1113–1121.

38. Chechneva, O.V. and W. Deng, Mitochondrial translocator protein (TSPO), astrocytes and neuroinflammation. Neural Regen Res, 2016. 11(7): p. 1056–7.

39. Daugherty, D.J., et al., The hGFAP-driven conditional TSPO knockout is protective in a mouse model of multiple sclerosis. Sci Rep, 2016. 6: p. 22556.

40. Sadeghian, M., et al., Mitochondrial dysfunction is an important cause of neurological deficits in an inflammatory model of multiple sclerosis. Sci Rep, 2016. 6: p. 33249.

41. Zhou, K., et al., Recent Advances of the NLRP3 Inflammasome in Central Nervous System Disorders. J Immunol Res, 2016. 2016: p. 9238290.

42. Fang, E.F., et al., *Mitophagy inhibits amyloid-*β *and tau pathology and reverses cognitive deficits in models of Alzheimer’s disease*. Nat Neurosci, 2019. 22(3): p. 401–412.

43. Frison, M., et al., The translocator protein (TSPO) is prodromal to mitophagy loss in neurotoxicity. Mol Psychiatry, 2021. 26(7): p. 2721–2739.

44. Gatliff, J., et al., TSPO interacts with VDAC1 and triggers a ROS-mediated inhibition of mitochondrial quality control. Autophagy, 2014. 10(12): p. 2279–96.

45. Issop, L., et al., Translocator Protein-Mediated Stabilization of Mitochondrial Architecture during Inflammation Stress in Colonic Cells. PLoS One, 2016. 11(4): p. e0152919.

46. Scaini, G., et al., TSPO upregulation in bipolar disorder and concomitant downregulation of mitophagic proteins and NLRP3 inflammasome activation. Neuropsychopharmacology, 2019. 44(7): p. 1291–1299.

47. Thai, P.N., et al., Cardiac-specific Conditional Knockout of the 18-kDa Mitochondrial Translocator Protein Protects from Pressure Overload Induced Heart Failure. Sci Rep, 2018. 8(1): p. 16213.

48. Gatliff, J., et al., A role for TSPO in mitochondrial Ca(2+) homeostasis and redox stress signaling. Cell Death Dis, 2017. 8(6): p. e2896.

49. Chen, K., et al., Optineurin inhibits NLRP3 inflammasome activation by enhancing mitophagy of renal tubular cells in diabetic nephropathy. Faseb j, 2019. 33(3): p. 4571–4585.

50. Gong, Z., et al., Mitochondrial dysfunction induces NLRP3 inflammasome activation during cerebral ischemia/reperfusion injury. J Neuroinflammation, 2018. 15(1): p. 242.

51. Zhang, N.-P., et al., Impaired mitophagy triggers NLRP3 inflammasome activation during the progression from nonalcoholic fatty liver to nonalcoholic steatohepatitis. Laboratory Investigation, 2019. 99(6): p. 749–763.

52. Huang, Y., W. Xu, and R. Zhou, NLRP3 inflammasome activation and cell death. Cell Mol Immunol, 2021. 18(9): p. 2114–2127.

53. Nutma, E., et al., Translocator protein is a marker of activated microglia in rodent models but not human neurodegenerative diseases. Nature Communications, 2023. 14(1): p. 5247.

