## Supplemental Methods for "Mitochondria drive microglial NRLP3 inflammasome activation via TSPO"

**Supplementary Methods**

**Protein extraction and subcellular fractionation**

Cells were resuspended in cell lysis buffer (50 mM Tris (Sigma, T6066) pH 8.0, 150 mM NaCl, 1% Triton-X (Sigma, T9284)) containing protease inhibitors (Roche, 04693132001) and kept on ice for 30 minutes. The volume was centrifuged at 10,000 RPM for 5 minutes at 4 ºC to allow for the removal of cellular debris.

Subcellular fractionation: The 3 subcellular compartments of interest (Cytosol, nucleus, and mitochondria) were separated by differential centrifugation. To isolate mitochondria, sucrose isotonic fractionation buffer (250 mM sucrose, 20 mM HEPES (pH 7.4), 10 mM KCI, 1.5 mM MgCl_2_, 1 mM EDTA, 1 mM EGTA) containing protease inhibitors are added to washed plates. The cells were scraped, and the suspension passed through the 26-gauge needle with 1 mL syringes 10-12 times before leaving on ice for 20 minutes. The suspension is centrifuged at 3000 RPM for 5 minutes. The subsequent pellet contains the nuclear fraction. The nuclear pellet is washed using fractionation buffer and passed through a 25-gauge needle and 1 mL syringe 10-12 times. This is followed by centrifugation at 3000 RPM for 10 minutes and the nuclear pellet is resuspended in nuclear buffer (standard lysis buffer containing 10% glycerol and 0.1% SDS). The remaining supernatant (after the initial centrifugation) is transferred and centrifuged at 8000 RPM to obtain the mitochondrial pellet. The supernatant, which consists of the cytosolic fraction, is stored at -20°C until required. The remaining pellet (mitochondrial fraction) is washed using isotonic buffer and centrifuged at 3000 RPM, for 5 minutes. Following this, the pellet is resuspended in cell lysis buffer and stored at -20 °C until required.

**Assessment of mitochondrial membrane potential (ΔΨ_m_)**

Tetramethylrhodamine, methyl ester (TMRM) (Thermo Fisher, T668) dye (100nM) was used to assess ΔΨ using a Leica SP5 to capture images (excitation at 548 nm and emission at 574 nm) after a 30 min incubation.

**Ca^2+^ signalling analysis**

To monitor cytoplasmic calcium between cell lines and with/without LPS treatment, we made use of a ratiometric dye, Fura-2 (Thermo Fishes, F-1221) which is bound to an acetoxymethyl (AM) ester increasing cell permeability. Cells were seeded 48 hours prior on 22 mm glass coverslips, LPS-treated cells were treated using LPS for 24 hours, and then all cells were washed using Hank’s Balanced Salt Solution (HBSS, Thermo fisher, 24020117) 24 hours later. The glass slides were loaded with 2.5 μM Fura-2 by diluting in HBSS and incubated at RT for 40 min. The cells were washed in HBSS 3 times, and the coverslips were mounted onto Attofluor Cell Chambers. The slides were imaged using a UV Nikon microscope where a field of cells was selected (>20 cells) by drawing ROIs. Continuous recording was set up (1 image per sec) to continuously capture at least 500 frames using a 40X objective. As the Fura-2 indicator is a ratiometric dye, the use of a 340/380 nm excitation ratio was necessary. After continuously recording for approx. 30 seconds/1 minute to get a basal intracellular reading, 100 μM of Adenosine triphosphate (ATP) was added to the slide. ATP acts on the purinergic receptors causing calcium uptake. The mobilization of Ca^2+^ results in a sharp increase in the intensity of Fura-2 fluorescence. When the signal returned to basal level values, the slides were treated with 5 μM ionomycin (Sigma, I3909), which increases cytosolic Ca^2+^, saturating the dye and resulting in a substantial increase in fluorescent intensity. Images were processed and analysed using the imaging software Andor iQ2 by demarcating cellular ROIs.

**Oxygen Consumption Rate (OCR) analysis**

Oxygen Consumption RateOCR analysis was performed using Agilent Seahorse 96 well XF Analyzer (Agilent Technologies) with XFe96 FluxPak (Agilent Technologies, 102416-100) according to manufacturer’s instructions. Additionally, XF DMEM medium, pH 7.4, 4 (Agilent Technologies, 103575-100), XF 1.0 M Glucose solution (Agilent Technologies, 103577-100), XF 100 mM Pyruvate solution (Agilent Technologies, 103578-100) and XF 200 mM Glutamine solution (Agilent Technologies, 103579-100) were purchased for use with the flux peak. 48 hours before assay WT and TSPO KO BV2 cells were seeded (3000) in triplicate for each treatment group on the 96-well dishes that accompany the fluxpak. 24 hours before the assay is performed a sensor cartridge is hydrated in Seahorse XF Calibrant (according to manufacturer’s instructions) at 37 C. Assay medium was prepared On the day of the assay, the medium was prepared (Seahorse XF DMEM, 1 mM pyruvate, 2 mM glutamine, and 10 mM glucose) and. Working solutions of the compounds were made using the assay medium (Concentrations used were 1.5 µM, 1 µM and 0.5 µM for Oligomycin, FCCP and Rotenone/antimycin A, respectively).

**Assessment of cellular viability**

MTT assay 3-(4,5-dimethylthiazol-2-yl)-2,5-diphenyltetrazolium bromide (MTT) is a colourimetric assay used to assess cell metabolic activity. Cells were seeded in triplicate in 24 well plates and treated (24h later) with either 100 ng/ml of LPS, 100 nM GE-180 or Vehicle. 24 h later, the cells were washed with PBS and incubated in RPMI-1640 (Gibco, Life Technologies), containing 0.5 mg/mL of MTT (Sigma Aldrich) followed by a 1hr incubation at 37˚C and 5% CO_2_ atmosphere. The supernatant was moved to 96-well plates, and the absorbance was measured using a Spectrophotometer (Tecan, UK) at 595 nm. Sample absorbance was normalized to control to give percentage absorbance.

Crystal violet is a triarylmethane dye which binds to proteins and DNA. Cells were seeded (in triplicate) and treated on 96 well plates then washed using PBS followed by the addition of L of 0.5% crystal violet staining solution. Following 20 minutes of incubation at RT, the plates are rinsed with H_2_O gently to remove all residual crystal violet. The plate is allowed to air-dry for 24 hours. 200 μL methanol was added to each well and incubated for 20 minutes at RT. Sample absorbance was read at 570nm with a Tecan Sunrise plate reader. The OD reading from wells containing complete media alone is subtracted from all readings to normalise for background.

**GE-180 binding assessment**

Ethical approval for human participants was obtained from the Research Ethics Board at McMaster University, Ontario, Canada. Reference number R.P. #12-3658. All procedures performed in studies involving human participants were by the ethical standards of the institutional and/or national research committee and with the 1964 Helsinki Declaration and its later amendments or comparable ethical standards. In total, 20 patients were considered, with 9 healthy patients and 11 with relapsing-remitting Multiple Sclerosis (RMS).

All processing was performed using PMOD software (PMOD Technologies v3.5, Zurich, Switzerland) using the PNeuro, PFusion and PView tools. A detailed description of the full methodology is described in supplementary materials

All processing was performed using PMOD software (PMOD Technologies v3.5, Zurich, Switzerland) using the PNeuro, PFusion and PView tools. The last 60-90 minutes of dynamic [18F]GE-180 PET scans were summed into static images using PView. Multiple ROIs were drawn from the start of the carotid to the end of the carotid by moving across the PET images, the multiple ROIs were linked to create a 3D carotid mask which would be applied later. The masks were visually inspected to ensure they encompassed the complete internal carotid arteries. The patient PET summation image was transferred into PNeuro where “standard tissue probability” was applied to the PET image in MR space, allowing segmentation of the image into either CSF, white matter or grey matter. The file was saved under the patient number and then opened in PFusion, here the previous segmentation processed for the patient image was separated to the range for white matter alone by segmentation and saved as a mask. Finally, the patient PET summation was opened in PView where either the mask alone was applied to each specific PET image from the patient cohorts, or the mask was applied in conjunction with the carotid masks, after which the SUV ratios were noted down for all patients.

For the MRI-based carotids, the contrast-enhance T1-weighted MRI images for the corresponding patients that were taken in conjunction with the PET imaging were opened in PView. On these images, Multiple ROIs were drawn around the whole carotid arteries and were then linked to create a 3D carotid mask which could be applied later. The masks were visually inspected as well as applied to the PET images to ensure they were aligned correctly to the carotids. Once they were complete the respective PET images for the patients were opened on PView and the binding of the white matter masks generated previously was assessed relative to that of the binding of the MRI-derived carotid masks.

130

Methodology to assess binding concerning periventricular white matter (PVWM) using an atlas-based approach implemented in FSL software (FMRIB Software library v.5.0.9, UK). Other resources used in the evaluation include MNI152 T1 whole head & brain templates (1mm), MNI152-defined vascular density map (open source) and JHU DTI-based WM Atlas (MNI152-defined & provided with FSL). The processing was performed by Ms Lovena Chedumbarum Pillay (Neuro Analyst, GE Healthcare). Subjects’ T1-w MRI images were pre-processed, undergoing reorientation, rotational alignment and cropping using FSL’s FSLREORIENT2STD and FLIRT libraries. Subject-specific [18F]GE-180 PET images were registered to their respective T1 weighted MRI images by applying rigid transform and mutual information cost function using FSL’s FLIRT. Brain tissue was extracted from each of the subject’s pre-processed MRI using FSL BET. Using FSL’s FLIRT and FNIRT libraries (affine and non-linear transforms, respectively) the forward spatial transform (T1 > MNI) for each subject’s T1-weighted brain MRI was estimated. An MNI152 spatially normalized contrast-enhanced T1 MRI was used to accurately identify the internal carotid artery from the MNI152-defined vascular density map. Using FSLROI the carotid arty was then delineated. A PVWM mask was generated from the JHU DTI-based white matter atlas (Provided with FSL). The inverse spatial transforms (MNI152 > T1) were estimated from the previously derived forward transforms, and applied to the MNI152-defined carotid artery VOI. These transforms were also applied to the PVWM masks, allowing the transformation of carotid VOI and PVWM masks into native patient space.

The mean standard uptake value (SUV) within the carotid artery VOI was estimated and normalized to the mean periventricular uptake, as well as the means for PVWM uptake (SUVrPVWM) alone, using FSLSTATS. By implementing the PET image carotid analysis pipeline, I was able to obtain masks (see representative images below) that could be applied to the PET and MR images to assess the binding of GE-180.
