## Supplemental Material for "Mitochondria drive microglial NRLP3 inflammasome activation via TSPO"

### Supplementary Material: Figure 1

a

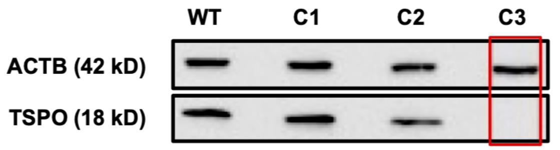


b

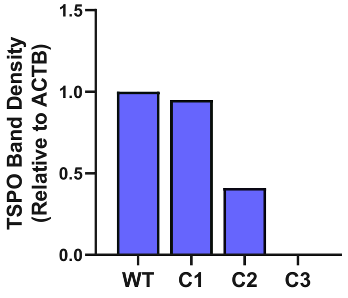


c

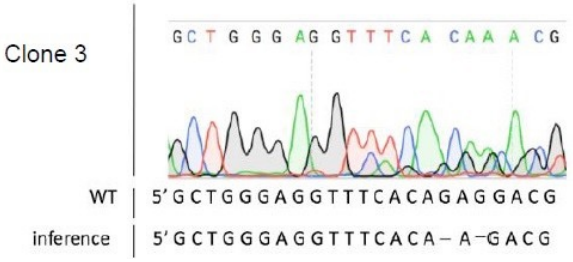


d

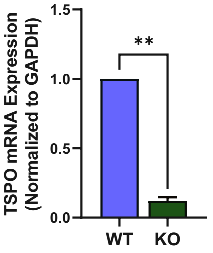

### Supplementary Material: Figure 2

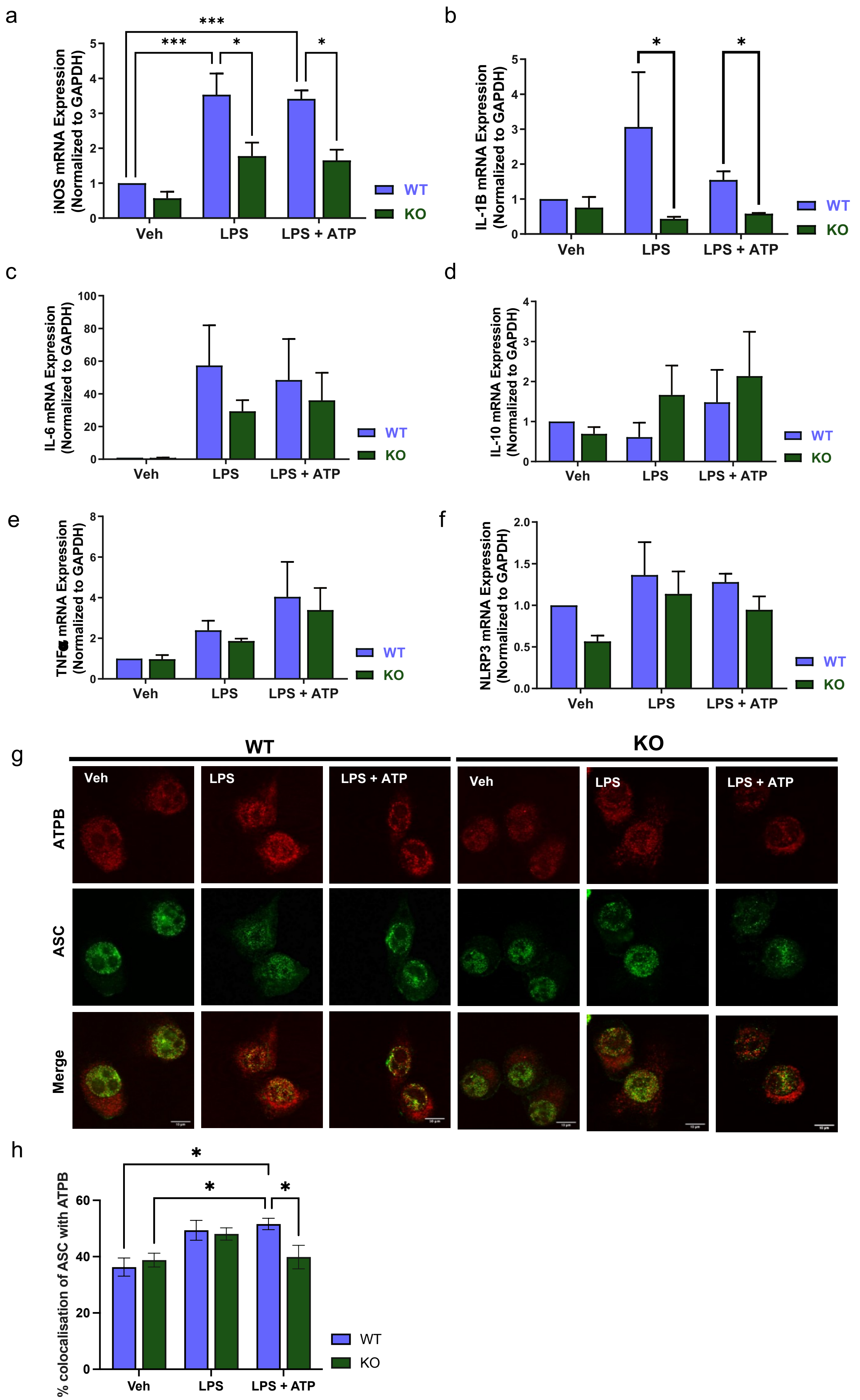

### Supplementary Material: Figure 3

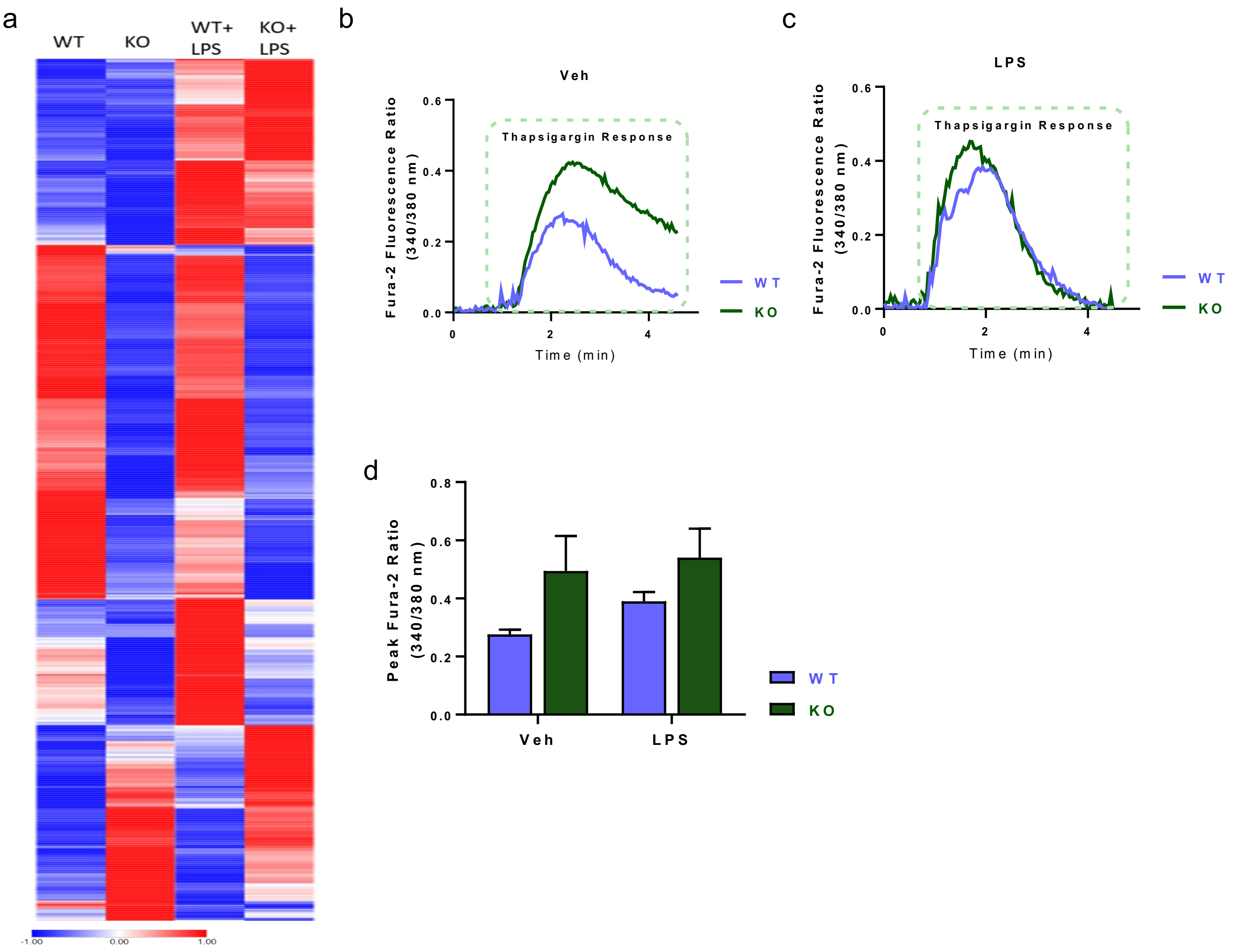

### Supplementary Material: Figure 4

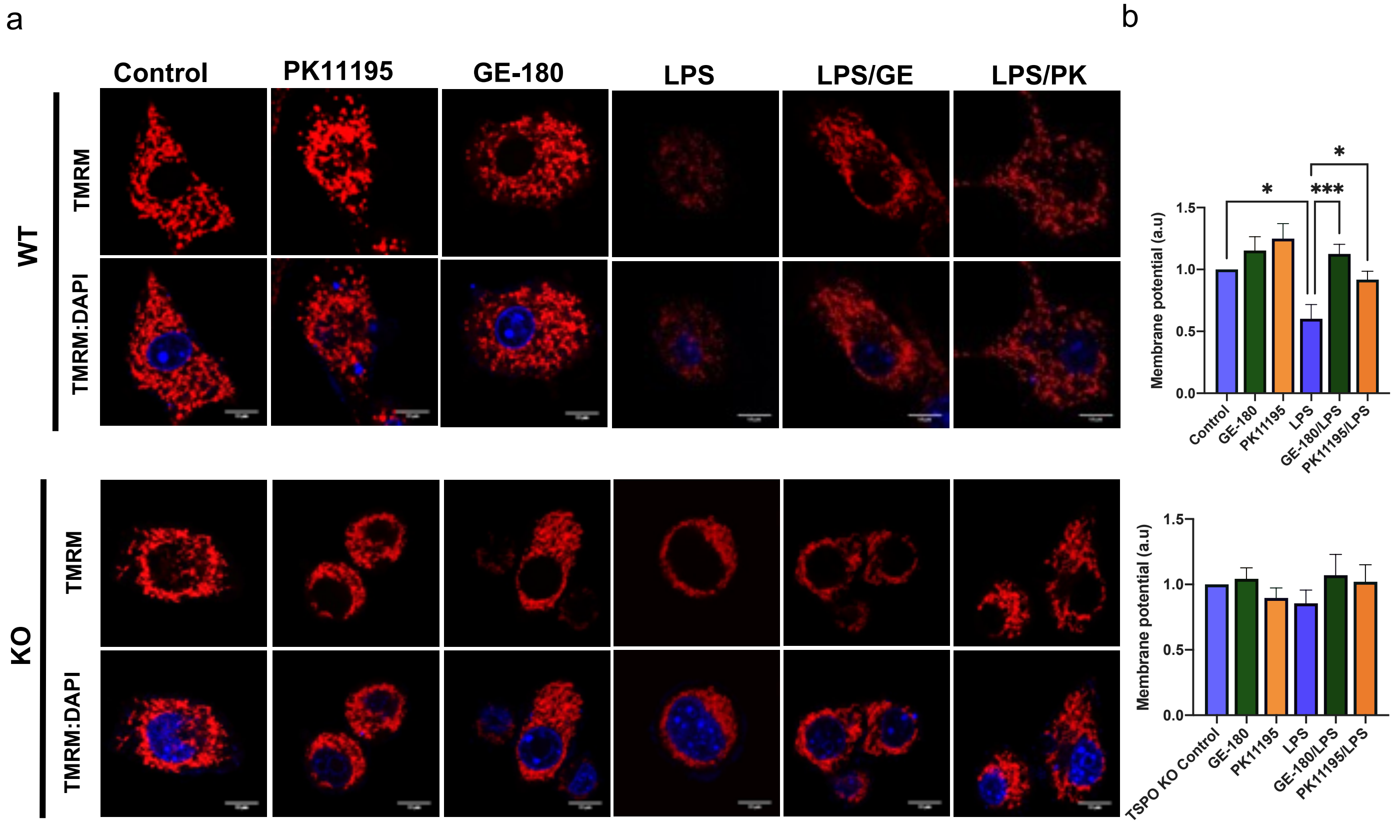

**Legends to Supplementary Material**

**Figure 1**

**a**, **b**) WT and CRISPR/Cas9-edited BV2 cell clones. (**a**) Western blot showing TSPO protein levels in WT BV2 cells and in selected negative (C1), heterozygous (C2) and homozygous (C3) TSPO KO clones; the blot confirms effective CRISPR/Cas9 gene editing-mediated ablation of TSPO in clone C3. (b) Quantification of TSPO protein levels in control and TSPO KO clones.
**c)** Representative chromatogram of TSPO KO cell showing a 2 base pair deletion in the TSPO gene, resulting in a frameshift mutation causing premature stop.
**d)** Histogram showing TSPO transcript levels in both WT and TSPO KO (clone 3) obtained via qPCR analysis, where the TSPO KO shows a significant decrease in TSPO copy number.

**Figure 2**

**a-f**) qPCR analysis of the expression levels of genes related to microglia activation in WT and TSPO KO BV2 cells treated with either vehicle control, LPS (100 ng/ml, 24 h) or LPS followed by ATP (2.5 mM, 30 min). Inducible nitric oxide synthase (iNOS) **(a)**, inflammatory cytokine IL1B **(b)**, inflammatory cytokine IL-6 **(c)**, anti-inflammatory cytokine IL-10 **(d)**, inflammatory cytokine TNF-a **(d)** and the NLRP3 protein **(e)**. Data are normalized to the control gene GAPDH and reported as fold changes in gene expression over control WT cells.
**g)** Representative co-immunocytochemical analyses of ASC and ATPB in WT and TSPO KO microglial cells untreated or exposed to LPS, LPS+ATP.
**h)**Histogram reporting the relative quantification

**Figure 3**

**a**) Heat maps of the RNA-se profiles in BV2 WT and BV2 TSPO KO cells, showing the overall difference in gene expression in resting conditions and when treated with LPS.
**b, c**) Thapsigargin-induced intracellular $Ca^{2+}$ rise (**c**) in untreated but not in LPS-treated cells (100 ng/mL, 24 h). This indicates that TSPO KO BV2 cells have a greater cytosolic $Ca^{2+}$ accumulation at rest. The summary of the amplitudes in bar chart (**d)** confirms the $Ca^{2+}$ response is higher in TSPO KO than in WT BV2 cells untreated, but that this response is attenuated under LPS treatment.

**Figure 4**

**a)** Representative images of TMRM accumulation in mitochondria in WT and TSPO KO microglial cells: untreated, exposed to LPS, exposed to TSPO ligands PK11195 and GE-180 or co-treated with a combination of the compounds.
**b)** Histograms reporting the relative quantification.
